# Identification of biomarkers and pathways of mouse embryonic fibroblasts with the dysfunction of mitochondrial DNA

**DOI:** 10.1101/2021.04.05.438453

**Authors:** Hanming Gu

## Abstract

Mitochondrial diseases are clinically heterogeneous which involve multiple systems such as organs that are highly dependent on metabolism. Dysfunction of mtDNA is the main cause of mitochondrial diseases that trigger inflammation and immune responses. Here, we aim to identify the biological function and pathways of MEFs with the dysfunction of mtDNA through deletion of YME1L. The gene expression profiles of GSE161735 dataset were originally created by the Illumina NovaSeq 6000 (Mus musculus) for gene biogenesis and function panel. The biological and functional pathways were analyzed by the Kyoto Encyclopedia of Genes and Genomes pathway (KEGG), Gene Ontology (GO), and Reactom visual map. KEGG and GO results showed the metabolism and immune responses were mostly affected by the loss of mtDNA. Moreover, we discovered several interacting genes including POLR2F, HIST1H2BJ, PPP1CC, HOXB4, ARG1, APITD1, BUB1B, POLR2K, HOXC4, and HOXB3 were involved in the regulation of metabolic or cancer diseases. Further, we predicted several regulators that had the ability to affect mitochondria during the dysfunction of mtDNA by L1000fwd analysis. Thus, this study provides further insights into the mechanism of mtDNA in metabolic diseases.

## Introduction

Mitochondria are critical regulators of metabolism in various cellular pathways, including fatty acid oxidation, oxidative phosphorylation, Krebs cycle, urea cycle, and gluconeogenesis^1^. Mitochondria also play a crucial role in other cellular processes such as amino acid metabolism, lipid metabolism, calcium homeostasis, and apoptosis^2, 3^. Mitochondrial diseases are complicated and involve mitochondrial and nuclear DNA mutations^4^. Generally, they cause the main defect in oxidative phosphorylation that involves ATP production, but lack of other enzymes such as Krebs enzymes also can lead to severe diseases^5-7^.

Mitochondrial DNA (mtDNA) encodes crucial proteins of the oxidative phosphorylation system^8^. This system contains an electron transport chain and ATP synthase, which makes mitochondrial respiration and ATP production^9^. Recently, it has been shown that mitochondrial DNA has various functions in immune responses^10^. Numerous evidence was identified for mtDNA regulating immune regulation^11^. Firstly, mtDNA is small and encodes 13 oxidative phosphorylation mRNAs, tRNAs, and ribosomal RNAs which are needed for mitochondrial matrix constitution^12^. Secondly, mtDNA copy number is controlled by cell-specific mechanisms and in response to various environmental stresses^13^. Thirdly, mtDNA contains unique bacterial nucleic acid sequences that make it more like foreign DNA^14^. Finally, mtDNA may exhibit persistent oxidative damage modifications or mutations due to its oxidative environment and DNA repair mechanisms^15^. Thus, mtDNA may have an essential role in innate immune response and inflammation.

In this study, we investigated the block of mtDNA in MEFs by using the protease YME1L. YME1L preserves nucleotide by supporting de novo nucleotide synthesis and limiting mitochondrial nucleotide transport and accumulation^16, 17^. We identified DEGs, candidate regulators, and the biological process in YME1L KO MEFs by performing a comprehensive bioinformatics analysis. The functional enrichment analysis and protein-protein interaction were utilized for discovering the significant gene nodes. These key genes and signaling pathways may be essential to therapeutic interventions of mitochondrial diseases.

## Methods

### Data resources

The GSE161735 dataset was downloaded from the GEO database (http://www.ncbi.nlm.nih.gov/geo/). It was produced by Illumina NovaSeq 6000 (Mus musculus) for cell biogenesis and function panel, Max Planck Institute for Biology of Ageing, Cologne, Germany. RNA-Seq analysis was performed using mouse embryonic fibroblasts (MEFs) with the knockout of YME1L gene (YME1L KO) as well as wild-type (WT) controls.

### Data acquisition and preprocessing

The GSE161735 dataset that contains gene expression related to gene functions from the MEFs with the knockout of YME1L gene (YME1L KO) was conducted by R script^18, 19^. We used a classical t-test to identify DEGs with P<0.01 and fold change ≥1.5 as being statistically significant.

### Gene functional analysis

Gene Ontology (GO) is a structured bioinformatics resource that includes three domains: biological processes (BP), cellular components (CC), and molecular functions (MF). Kyoto Encyclopedia of Genes and Genomes (KEGG) database is the pathway tool that integrates functional information and the known and unknown biological pathways. GO and KEGG pathways were conducted by utilizing the Database for Annotation, Visualization, and Integrated Discovery (DAVID) (http://david.ncifcrf.gov/) and Reactome (https://reactome.org/)^20^. P<0.05 was considered statistically significant.

### Module analysis

We study the connected regions in protein-protein interaction (PPI) networks using the Molecular Complex Detection (MCODE) of Cytoscape software^21, 22^. The significant clusters were selected from the constructed PPI network using MCODE and String database (https://string-db.org/). The pathway enrichment analyses were performed by using Reactome, and P<0.05 was used as the cutoff criterion.

### Reactome analysis

Reactome pathway (https://reactome.org/) was conducted to obtain the visualization analysis of potential pathways. P<0.05 was considered statistically significant.

## Results

Identification of DEGs of Yme1l KO MEFs in comparison to those of WT MEFs Mitochondrial DNA (mtDNA) elicits a type I interferon response, but the mechanism of mtDNA release remains unknown. The mitochondrial protease YME1L maintains nucleotide pools by supporting de novo nucleotide synthesis and the accumulation of mtDNA. Transcript profiles of WT and Yme1l KO MEFs were analyzed by differentially expressed genes (DEGs) from the Max Planck Institute for Biology of Ageing, Germany. A total of 3495 genes were identified to be differentially expressed in Yme1l KO MEFs with the threshold of P<0.05. The top 10 up-and down-regulated genes are listed in Table 1.

**Table 1.**
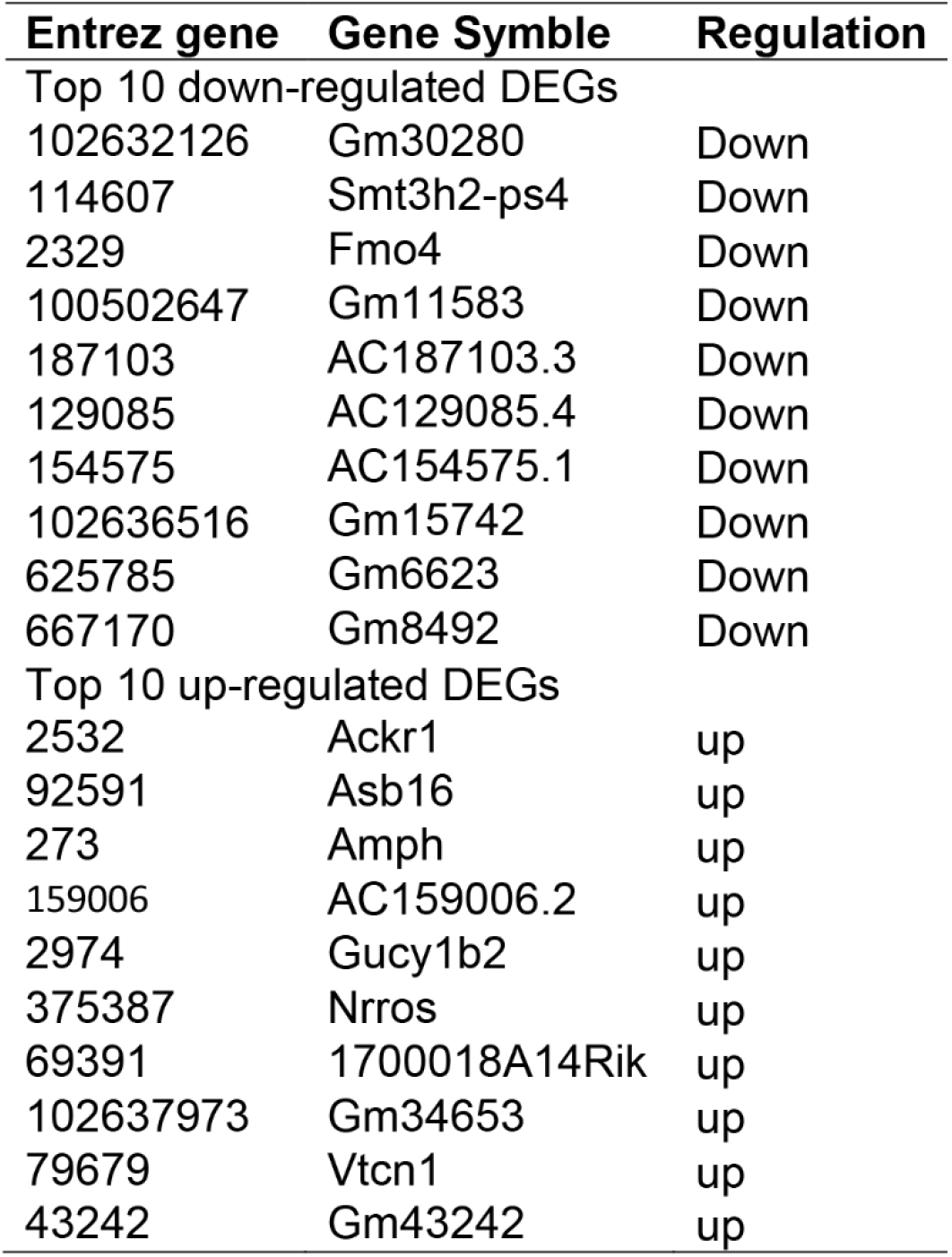

### KEGG analysis of DEGs of Yme1l KO MEFs in comparison to those of WT MEFs

To identify the biological functions and pathways of DEGs of Yme1l KO MEFs, we conducted a KEGG pathway enrichment analysis (Supplemental Table S1). KEGG pathway is a dataset collection for analyzing molecular interaction and relation networks. Our study showed the top ten enriched KEGG pathways including “Metabolic pathways”, “MAPK signaling pathway”, “Herpes simplex infection”, “Parkinson’s disease”, “Huntington’s disease”, “Oxidative phosphorylation”, “Autoimmune thyroid disease”, “Leishmaniasis”, “Metabolism of xenobiotics by cytochrome P450”, and “Antigen processing and presentation” (Figure 1).

**Figure 1.**
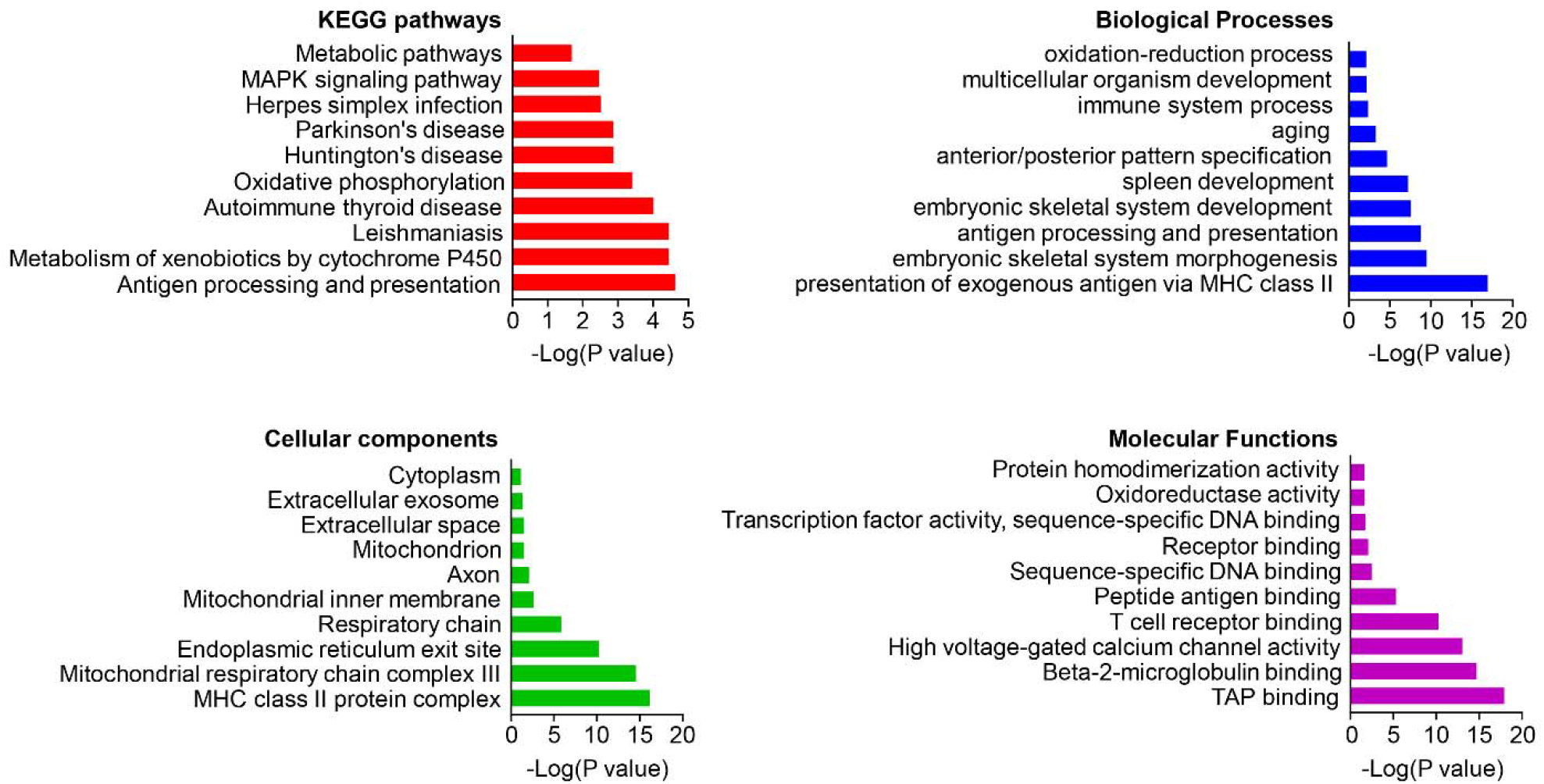
The KEGG pathways, biological process, cellular component, and molecular function terms enriched by the DEGs. DEGs =differentially expressed genes, KEGG = Kyoto Encyclopedia of Genes and Genomes.

### GO analysis of DEGs from Yme1l KO MEFs

Gene Ontology (GO) analysis is a useful tool for sorting genes, which contains cellular components (CC), molecular functions (MF), and biological processes (BP). We identified the top ten cellular components including “cytoplasm”, “extracellular exosome”, “extracellular space”, “mitochondrion”, “axon”, “mitochondrial inner membrane”, “respiratory chain”, “endoplasmic reticulum exit site”, “mitochondrial respiratory chain complex III”, and “MHC class II protein complex” (Figure 1). We also identified the top ten biological processes: “oxidation-reduction process”, “multicellular organism development”, “immune system process”, “aging”, “anterior/posterior pattern specification”, “spleen development”, “embryonic skeletal system development”, “antigen processing and presentation”, “embryonic skeletal system morphogenesis”, and “antigen processing and presentation of exogenous peptide antigen via MHC class II” (Figure 1). We then identified the top ten molecular functions: “protein homodimerization activity”, “oxidoreductase activity”, “transcription factor activity, sequence-specific DNA binding”, “receptor binding”, “sequence-specific DNA binding”, “peptide antigen binding”, “T cell receptor binding”, “high voltage-gated calcium channel activity”, “beta-2-microglobulin binding”, and “TAP binding” (Figure 1 and Supplemental Table S1).

### PPI network and Module analysis

PPI networks were created to analyze the relationships of DGEs at the protein level. The criterion of combined score >0.7 was conducted and the PPI network was constructed by using 301 nodes and 627 interactions. Among these nodes and edges, the top ten genes with the highest scores are shown in Table 2. The significant modules of DEGs of Yme1l KO MEFs in comparison to those of WT MEFs were selected to show the functional annotation (Figure 2).

**Table 2.**
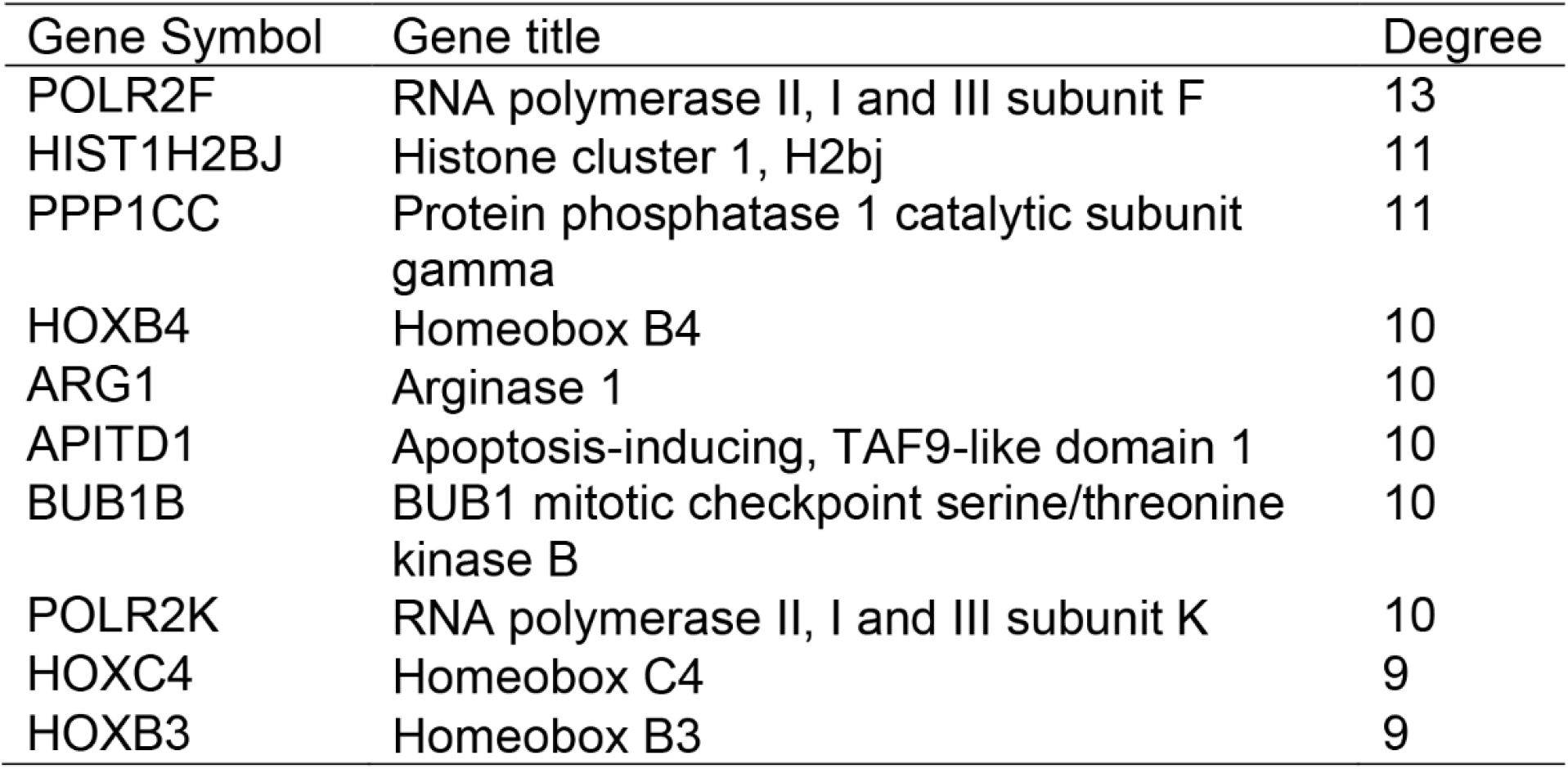
Top ten genes demonstrated by connectivity degree in the PPI network.

**Figure 2.**
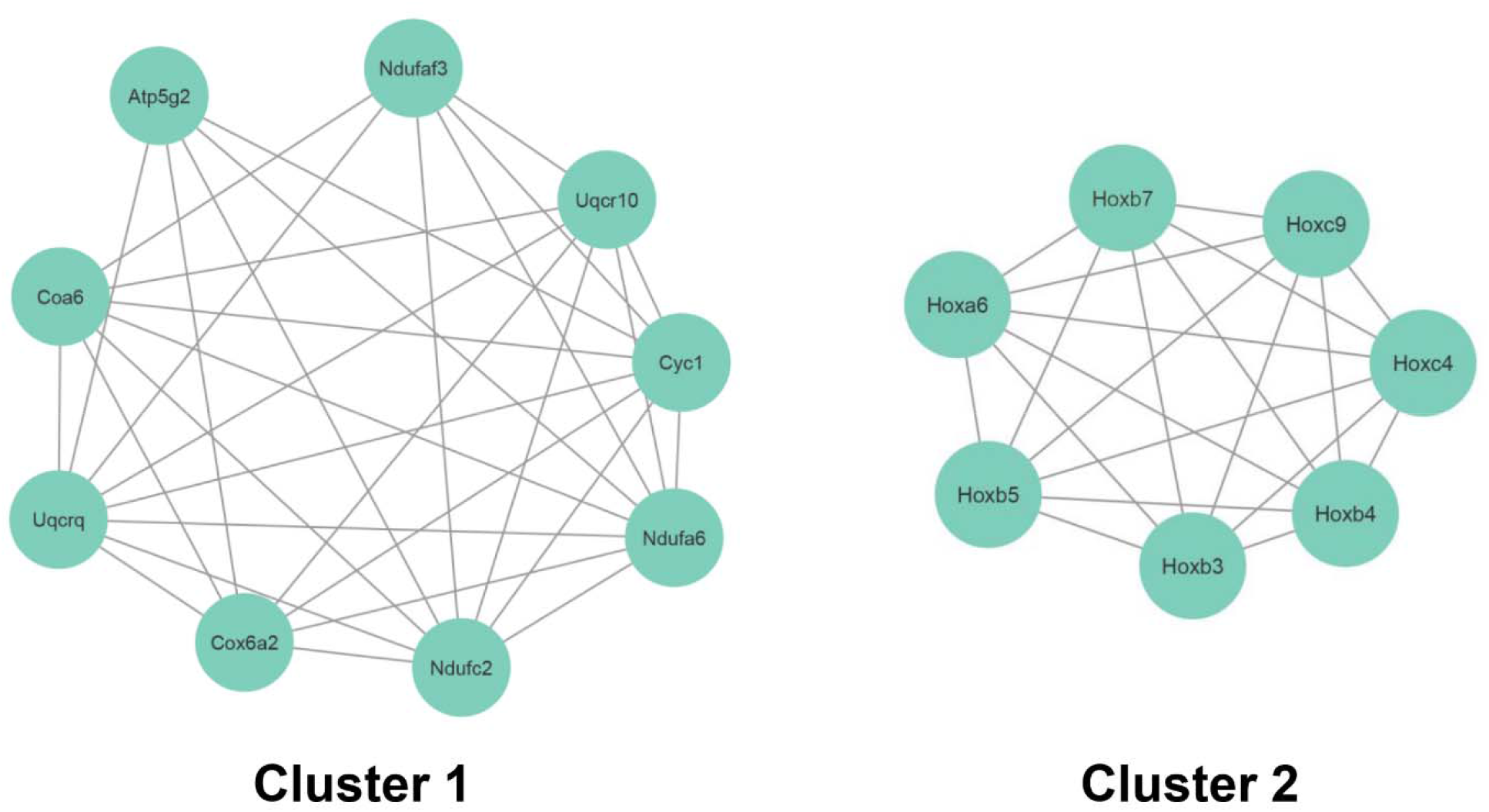
Top two modules from the PPI network.

### Reactome Pathway analysis

To further understand the potential functions of DEGs, we also identified the signaling pathways by using Reactome Pathway Database. The top ten signaling pathways: “Response of EIF2AK1 (HRI) to heme deficiency”, “Activation of anterior HOX genes in hindbrain development during early embryogenesis”, “Activation of HOX genes during differentiation”, “ATF4 activates genes in response to endoplasmic reticulum stress”, “PERK regulates gene expression”, “Defective Base Excision Repair Associated with OGG1”, “Response of EIF2AK4 (GCN2) to amino acid deficiency”, “Respiratory electron transport”, “EPHA-mediated growth cone collapse”, and “Rhesus blood group biosynthesis” (Supplemental Table S2). We then constructed the visual reaction map according to the signaling pathways (Figure 3).

**Figure 3.**
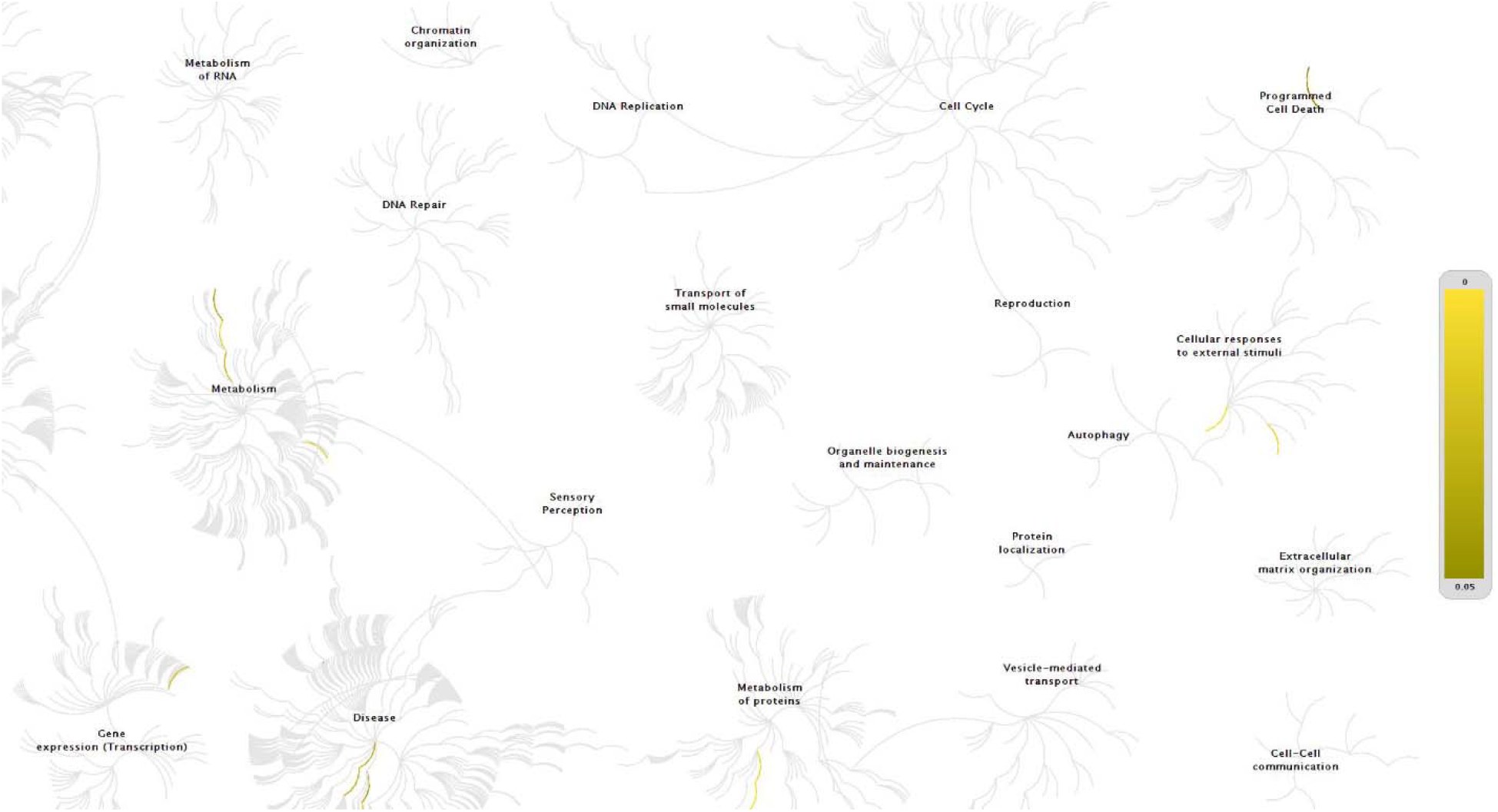
The Reactom pathways. The top significantly changed genes obtained from the GSE161735 dataset (P <0.01). The yellow color represents the most relevant signaling pathways.

### Potential regulators analysis

To further know the potential regulators, we use the L1000 fireworks display system that can predict bioactive molecules. The system indicated the potential pathways that may be inhibited. We selected the top ten molecules according to the DEGs and inhibitor map: “vincristine”, “BRD-K89059493”, “VU-0410183-2”, “indirubin”, “heliomycin”, “AT-SUMO-1”, “PD-98059”, “tosyl-phenylalanyl-chloromethyl-ketone”, “loteprednol”, and “BRD-K51556300” (Figure 4 and Supplemental Table S3).

**Figure 4.**
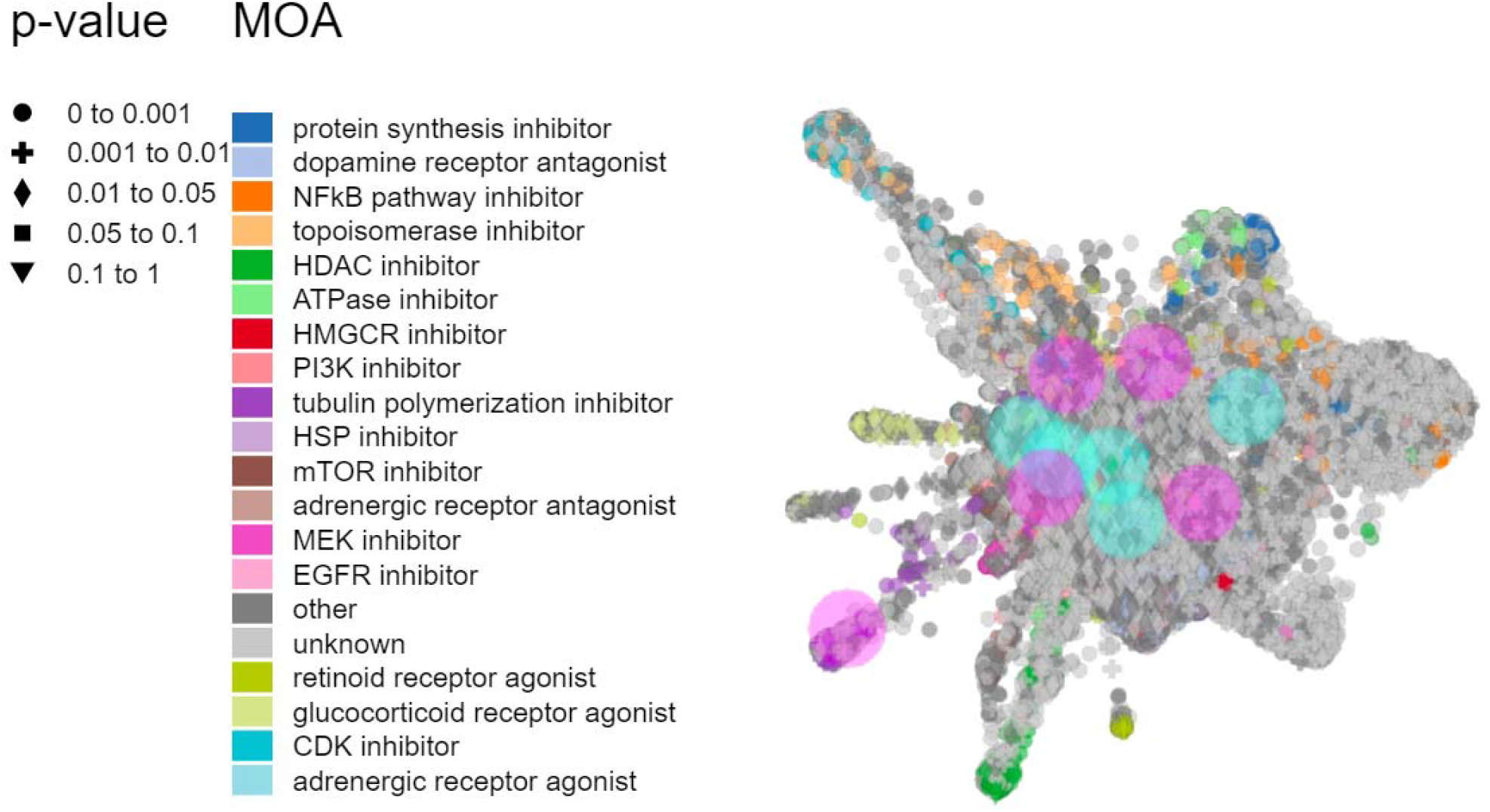
Inhibitor predictions by L1000FDW. The significantly changed genes obtained from the GSE161735 dataset. Dots are the Mode of Action (MOA) of the respective drug.

## Discussion

Mitochondria are key scaffolds of cellular metabolism and signaling, but they are also being proved as critical participants in immune responses to pathogens and cellular damage^23^. Mitochondria are not only necessary for antiviral signaling but also the essential sources of endogenous DAMPs^11^. mtDNA triggers pro-inflammatory processes and type I IFN responses owing to its unique structural features and heightened susceptibility to oxidative damage^24^. Thus, future work to disclose the mechanism of mtDNA will have broad implications for knowing mitochondrial diseases, perhaps leading to new avenues for therapy to improve human health.

To understand the effects of mtDNA, we analyzed the gene expressions related to YME1L KO MEFs in this study. By analyzing the DEGs, ten proteins were identified according to the PPI network analysis, which may be important in mitochondrial diseases or other diseases. For example, POLR2F is related to the periventricular white matter hyperintensities (WMH; PVWMH)^25^. HIST1H2BJ is identified for predicting the prognosis of cervical cancer patients^26^. PPP1CC is required for SLFN11 to cleave tRNAs and further to inhibit the translation^27^. G protein, GPCR, and related proteins are involved in widely physiological and pathophysiological processes^28-33^. PPP1CC plays an important role in regulating the GPCR signaling pathways^34^. HOXB4 can induce the expansion of hematopoietic and promote the malignant progression of ovarian cancer^35, 36^. Arg1 is related to the tumor-associated macrophage and inflammation resolution^37^. Circadian gene clocks regulate numerous physiological processes such as metabolism, immune, and aging^38-41^. A recent finding is that Arg1 is up-regulated in the Rev-erb KO mice^42^. APITD1 contains a p53-binding domain, which indicates low expression in neuroblastoma tumors^43^. BUB1B is positively correlated with cancer progression and promotes hepatocellular carcinoma through triggering the mTORC1 signaling pathway^44^. POLR2K involves prostate cancer progression and lethal disease^45^. HOXC4 controls prostate cancer cells via regulating genes and binding sites^46^. HBOX3 is associated with the progression of prostate cancer^47^. Thus, dysfunction of mtDNA may cause potential diseases not only related to mitochondria but other kinds of diseases as well.

KEGG and GO analyses indicated that metabolism and immune were the main pathological processes after blocking the mtDNA. The processes of “Metabolic pathways”, “Oxidative phosphorylation”, “Metabolism of xenobiotics by cytochrome P450” and “oxidation-reduction process” in KEGG and GO were significantly changed. Since mitochondria are the main organelle in cells for energy production, the loss of the function of mtDNA will probably cause metabolic dysfunctions. We also found “Autoimmune thyroid disease”, “Antigen processing and presentation” and “immune system process” were involved in the process of mtDNA blocking. Similarly, mtDNA acts as a stress that promotes the antiviral innate immune response^48^. mtDNA activates the innate immune cGAS/STING pathway in the absence of apoptosis, which causes the dying cells to secrete type I interferon^49^. mtDNA also involves the NF-κB signaling pathway that widely regulates the inflammatory factors^50-52^. It was reported that the dysfunction of mitochondria leads to bone defect and bone diseases^53^. Here, we also found the block of mtDNA can affect embryonic skeletal system development and its morphogenesis.

In summary, we discovered the biological pathways in Yme1lKO MEFs by analyzing gene functions. Metabolism and immune response are the mainly triggered pathways during the loss of mtDNA. Future studies will focus on the development of the potential regulators of mitochondrial diseases based on our study. Our study provides further insights into the dysfunction of mtDNA, which may facilitate drug development.

## Notes

### Competing Interest Statement

The authors have declared no competing interest.

